# Content of and interactions between repetitive elements in programmatically eliminated chromosomes of the sea lamprey (*Petromyzon marinus*)

**DOI:** 10.1101/567370

**Authors:** Vladimir A. Timoshevskiy, Nataliya Y. Timoshevskaya, Jeramiah J. Smith

## Abstract

The sea lamprey (*Petromyzon marinus*) is one of few vertebrate species that is known to reproducibly eliminate large fractions of its genome during normal embryonic development. In lamprey, elimination events are initiated at the 6^th^ embryonic cleavage and result in the loss of ∼20% of an embryo’s genomic DNA from essentially all somatic cell lineages (these same sequences are retained in the germline). This germline-specific DNA is lost in the form of large fragments, including entire chromosomes, and available evidence suggests that DNA elimination acts as a permanent silencing mechanism that prevents the somatic expression of a specific subset of “germline” genes. However, reconstruction of eliminated regions has proven challenging due to the complexity of the lamprey karyotype (84 small pairs of somatic chromosomes and ∼100 pairs of germline chromosomes), the exceedingly high repeat content of the genome and even higher repeat content of eliminated fragments.

We applied an integrative approach aimed at further characterization of the large-scale structure of eliminated segments, including: 1) *in silico* identification of germline-enriched repeats; mapping the chromosomal location of specific repetitive sequences in germline metaphases, and verification of repeat specificity to eliminated chromosomes by 3D DNA/DNA-hybridization to embryonic lagging anaphases. Our integrative approach resulted in the discovery of multiple highly abundant repetitive elements that are found exclusively on the eliminated (germline-specific) chromosomes which in turn permitted the identification of 12 individual chromosomes that are programmatically eliminated during early embryogenesis. The fidelity of germline-specific repetitive elements and their distinctive patterning in elimination anaphases are taken as evidence that these sequences might contribute to the specific targeting of chromosomes for elimination and possibly in molecular interactions that mediate their decelerated poleward movement in chromosome elimination anaphases, isolation from the primary nuclei and eventual degradation.

**AUTHOR SUMMARY:** Epigenetic silencing methods provide a means of precisely restricting gene expression while maintaining the integrity of the genomic template that encodes this information, and are employed by diverse species throughout the tree of life. Programmed genome rearrangement (PGR) represents a parallel approach that maintains genome integrity across generations but alters the genomes of cells within an organism. To better resolve elimination events that take place during PGR in the sea lamprey (one of few vertebrate species known to undergo large scale PGR) we sought to identify sequences that define specific eliminated chromosomes. Using computational predictions and cytogenetic validation, we identified six new repetitive elements that are restricted to the eliminated chromosomes and permit the identification of twelve distinct eliminated chromosomes. Analysis of these repeats in meiotic testes and in embryos sampled during the process of elimination shows that these repeats localize to specific subcellular regions, and suggest a potential role of these repetitive elements in targeting chromosomes for silencing via elimination.

## INTRODUCTION

The sea lamprey (*Petromyzon marinus*) possesses a distinctive mechanism of differentiating somatic and germline lineages that is achieved by discarding large portions of the genome during early stages of development [1, 2]. Similar largescale changes in genome content and structure have been described in several groups of phylogenetically diverse organisms [3] including: ciliates [4-6], nematodes [7, 8], sciarid flies (reviewed in [9]), copepods [10], chironomids [11], several hagfish species [12-15], songbirds [16, 17], and at least two lamprey species [18, 19]. These changes in genome content/structure are generally known as programmed genome rearrangement (PGR). The diversity of subcellular events associated with DNA elimination and the patchy taxonomic distribution of PGR reflect the repeated evolution of independent mechanisms regulating the reproducible targeting and processing of segments that are slated for elimination in somatic cell lineages.

Despite the diverse origins of PGR within Metazoan lineages, the loss of germline-specific segments shares common features across distantly related taxa. In most taxa elimination appears to largely play out during the progression of specific early embryonic anaphases, wherein eliminated chromatin exhibits differential motion relative to somatically retained chromatin (*e.g.* as observed in nematodes, copepods, chironomids, sciarid flies and lampreys). This general pattern can be divided into nonexclusive two categories: chromosome elimination per se that involves the removal of whole intact chromosomes (hagfish [20], sciarid flies [9] and song birds [16, 17, 21]) and chromatin diminution which includes steps of excision and rejoining, or telomere restoration (roundworms [22, 23], copepods [24, 25], chironomids [26] and some hagfish [27]).

Programmed deletions have been reported to target chromosomal segments containing highly repetitive DNA which are often packaged into transcriptionally repressive heterochromatin prior to elimination [8, 27-30]. However, in Taiwanese hagfish *Paramixine sheni* it has been reported that both C-band positive chromatin (presumptive heterochromatin) and C-band negative chromatin (presumptive euchromatin) are eliminated from somatic tissues [27]. More recent genome sequencing studies have revealed that programmed deletions result in the loss of hundreds of protein-coding genes in *Ascaris* [8] and the sea lamprey [2, 31], many of which are thought to function in the proliferation, development and differentiation of germ cells. The presence of both protein coding genes with presumptively critical functions and numerous high copy elements raises the questions as to whether repetitive elements themselves are truly junk targeted for elimination, passive passengers that are simply being carried along for the ride, or perhaps functionally relevant sequences that actively participate in the process of elimination.

In the present work, we focused on cytogenetically recognizable aspects of PGR in the sea lamprey. The sea lamprey’s somatic karyotype consists of 168 (n=84) small dot like chromosomes [18, 32, 33]. Its germline karyotype was previously estimated to consist of ∼99 small chromosomes, indicating, that lampreys might eliminate up to 15 entire chromosomes during PGR [18, 34]. However, the complex morphology of germline metaphase spreads and the presence of numerous small chromosomes have thus far prevented a precise description of the germline karyotype. Moreover, it has remained unclear the degree to which differences in chromosome number arise from wholescale loss of chromosomes *vs.* breakage and joining of remodeled segments.

To further characterize the content and distribution of germline-specific repetitive DNA in the sea lamprey, we performed computational analyses to identify several candidate repetitive elements that appeared to be highly enriched in germline and cytogenetic analyses to more precisely define their locations within both the germline-specific chromosomes during meiosis and within structured lagging chromatin that arises during the execution of PGR. These analyses provide a means of individually identifying chromosomes that are targeted by PGR, provide strong evidence that most (if not all) elimination events are achieved by the wholesale elimination of chromosomes, and identify elements that mark specific subregions of elimination anaphases. Notably, these repetitive elements include one highly-specific and abundant element (*Germ*2) that appears to mark a persistent zone of interaction between lagging anaphase chromatids that spans their original metaphase plane.

## RESULTS AND DISCUSSION

### Comparative hybridization indicates the presence of germline-specific repeats

To assess whether eliminated chromatin was likely to be enriched with germline-specific repeats, we performed comparative hybridization of repetitive DNAs (C_0_t2 fractions) from somatic (liver) and germline (testes) genomic DNA within intact embryos [19, 35]. Hybridizations were carried out on embryos that were actively undergoing elimination at the time of fixation (1.5-2 days post fertilization: dpf). Fluorescence intensity was measured using unprocessed images of cells containing micronuclei (Fig 1, A) and mean integrated fluorescence density of primary nuclei was used as background fluorescence respective to micronuclei. Micronuclei were found to be highly enriched in germline-derived repetitive DNA (p<0.0001, DF = 32) (Fig 1, B). A similar enrichment was also observed in the analysis of ∼30 eliminating anaphases: Cy3-labeled germline repeats exhibited sufficiently higher fluorescence intensity within lagging chromatin than FITC-labeled somatic repeats (Fig 1, C and S1 File). In eliminated chromatin, somatically retained repeats were observed to hybridize primarily with pericentromeric and peritelomeric regions forming dot-shaped signals and were largely absent from the internal regions of lagging chromosomes that hybridize with germline repetitive DNA (S1 File). This is interpreted as evidence that germline-specific and somatically-retained chromosomes share a similar complement of pericentromeric repeats and, in conjunction with the observation that the centromeres of eliminated chromosomes exhibit poleward motion during elimination anaphases, indicates that both eliminated and retained centromeres retain a capacity to form functional kinetochores.

**Fig 1.**
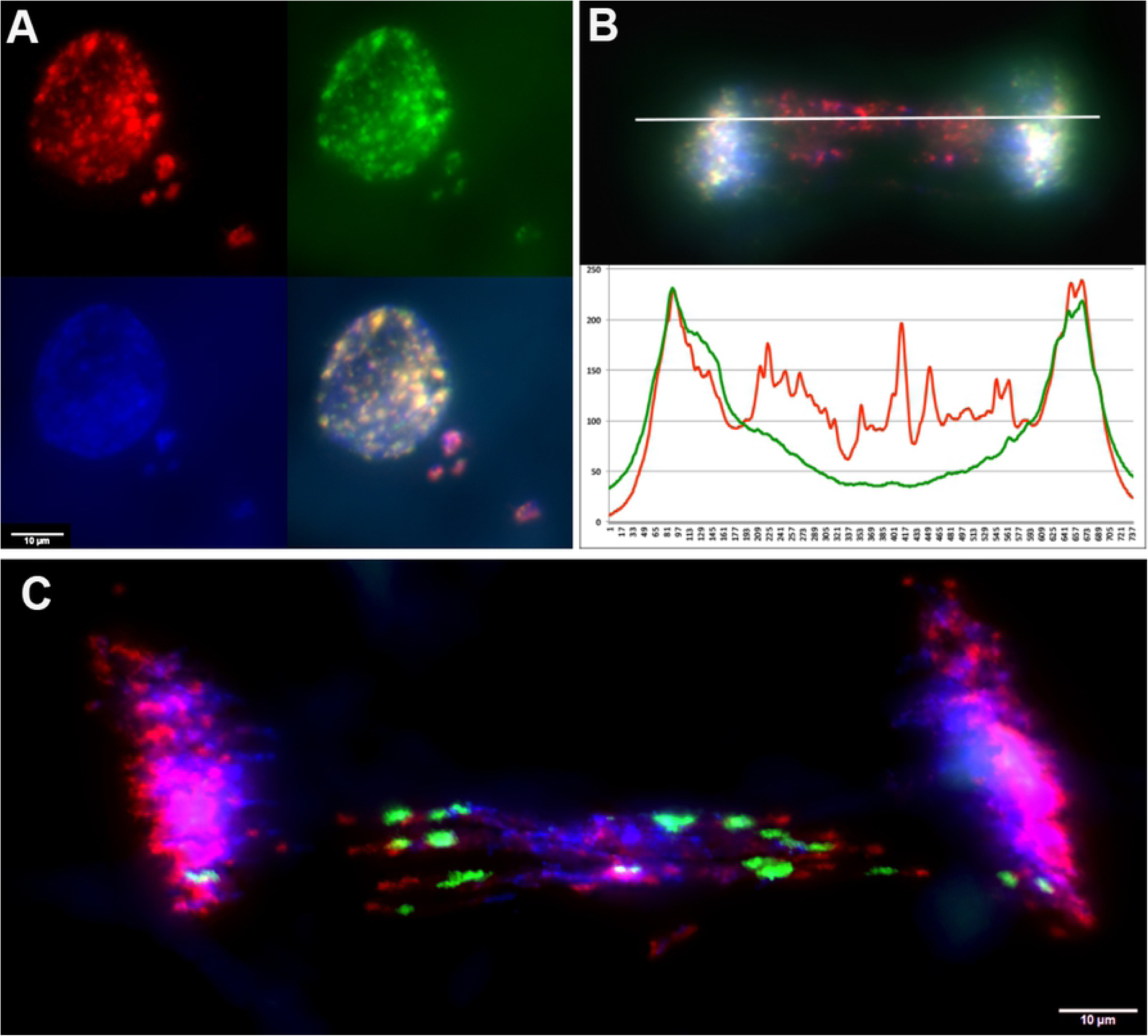
Repetitive DNA *in situ* hybridization in eliminating cells of 1.5 dpf sea lamprey embryos. (A) An example of competitive hybridization repetitive DNA fraction from testes (red) and liver (green) in a cell containing micronuclei. (B) Hybridization of C_0_t2 DNA-probes from somatic (green) and germline (red) to a lagging anaphase and a corresponding fluorescence intensity profile (measurement plane is marked by a horizontal line). (C) Hybridization of C_0_t1 (red) and *Germ*1 (green) to a lagging anaphase.

Previous studies have shown that a majority of micronuclei in 1-3 dpf sea lamprey embryos contain the germline-enriched repeat *Germ1* [2]. In order to determine whether germline C_0_t2 hybridization patterns could be explained simply by the presence of *Germ*1, we co-hybridized a *Germ*1 specific probe with labeled C_0_t1 DNA to elimination anaphases (Fig 1, C). Owing to its highly repetitive nature, labeled C_0_t1 DNA hybridizes to centromeric sequences and a small subset of subtelomeric repeats. These hybridizations reveal that *Germ*1 localizes to discreet regions within eliminated chromosomes, primarily near the centromeric ends of these largely acrocentric chromosomes, with the remaining portions of eliminated chromosomes being largely devoid of *Germ1*. Consistent with these observations, *Germ*1 probes were also found to hybridize to a subset of micronuclei (68% in D2 embryos and 66% in D2.5 embryos), whereas germline C_0_t2 DNA (repeats) hybridized to all micronuclei in D2 and D2.5 dpf embryos. These observations were taken as evidence that repetitive elements in addition to *Germ*1 were likely to be enriched within germline-specific chromatin, present in regions outside of the defined *Germ*1 clusters and, in some cases, fragmented from *Germ*1-positive regions prior to packaging as micronuclei.

### Computational identification of germline specific repeats

Consistent with the above findings, previous computational analyses of the repetitive content of germline and somatic DNA indicated that there are likely a large number of distinct repetitive elements that are unique to the germline [31]. To generate more precise consensus assemblies of individual repeat families and relative copy number estimates for each family, we performed a *de novo* assembly of repetitive elements that was seeded from a complete list of 31-mers that were abundant in sperm and/or blood reads. This assembly yielded a total of 130,632 consensus repetitive sequences. These sequences were merged with repeats that were identified within genomic scaffolds via RepeatModeler and an updated sequence of the *Germ*1 [36], yielding a total of 119,842 model repeats. Copy number estimates for consensus repeats were generated by remapping sequencing data from sperm and blood to obtain separate metrics of sequence coverage. A majority of repetitive sequences were abundant in both sperm and blood, however, the distribution of coverage ratios contains a notable tail corresponding to germline enriched sequences (Fig 2 and S1 Table).

**Fig 2.**
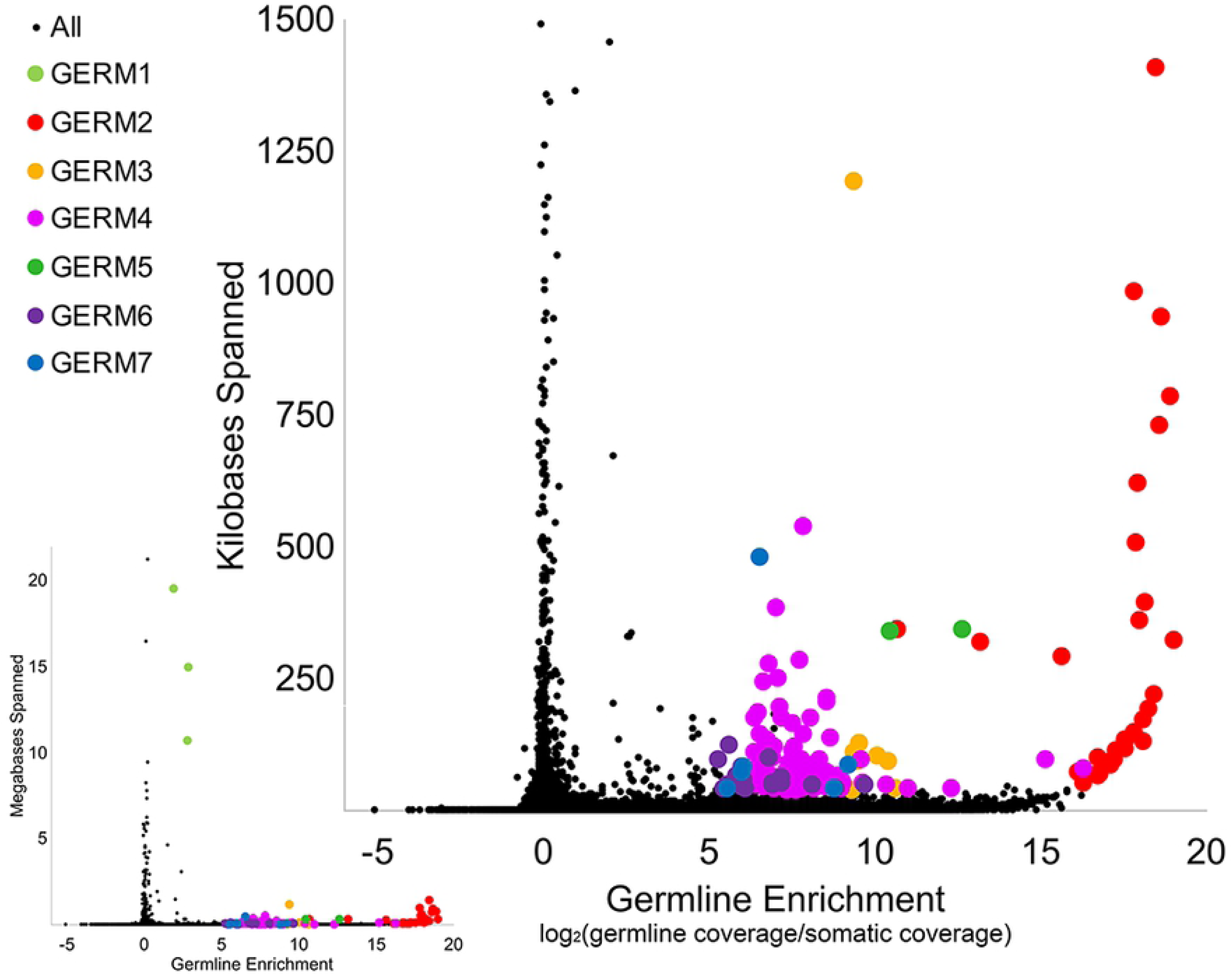
Estimates of the total genomic span and germline enrichment of repetitive sequences in the lamprey genome. The plot on the lower left shows enrichment values for the entire collection of reconstructed repetitive elements, including *Germ1*. The larger plot on the upper right focuses on repeats that span <1.5 megabases. Sequences belonging to repeat classes selected for hybridization are highlighted by colors corresponding to their repeat class.

All predicted high-copy elements with enrichment scores [log2(standardized sperm coverage/blood coverage)] exceeding 5 and an estimated span exceeding 40 kb, when summed across all copies, were extracted for downstream analysis (S2 Table). Subsequent inspection of these 171 predicted elements revealed similarities among subgroups of repeats, and semi-automated clustering revealed that these high-copy repeats could be grouped into 20 distinct clusters, *i.e.* repeat families. The representatives of 6 clusters, with a combined span size of more than 500Kb, were designated *Germ*2 – 7 (Fig 2 and S2 Table).

Examination of the sequences of *Germ*2 – 7 and genomics scaffolds (PIZI00000000.1) containing these repeats revealed that all of these high-copy germline-specific repeats occur as tandem arrays. Each repetitive element appears to contain a short (13-57bp) somewhat conserved core sequence (S3 Table). Tandem arrays of these core sequences are frequently disrupted by small insertions or deletions and, at larger scales, cassettes of tandem repeats are further duplicated as inverted repeats.

Arbitrarily chosen representatives of each cluster were selected for primer design for PCR validation. Amplicons generated from these primers yielded a continuous range of fragment sizes (smear) as would be expected for primers designed within tandem repeats, and relative specificity to the germline (S2 File). These same amplicons were used to generate probes (*Germ* 2 – 7) for subsequent FISH analyses.

### Defining the Chromosomal Localization of Germline-Restricted Repeats

The lamprey karyotype is characterized by a large number of small chromosomes, which presents significant challenges in the identification and characterization of individual homologs. Because repeats were predicted to be highly enriched in germline, we performed hybridizations against meiotic metaphases (spermatogenesis metaphase I) in order to characterize the distribution and location of these repetitive elements across lamprey chromosomes. Germline-specific chromosomes were identified on the basis of hybridization to a previously described laser-capture painting probe: LC [19]. Additionally, we used two probes, a telomere specific repeat (PNA probe) and centromere markers (labeled C_0_t1 fraction of genomic DNA from somatic lineage) to aid defining the bounds of individual chromosomes (S3 File). These hybridizations yielded counts of 84 somatically retained and 12 germline-specific chromosomes. In meiotic spreads the LC probe hybridized with several entire chromosomes and a portion of one somatically-retained chromosome that has been previously shown to hybridize to *Germ*1 in somatic metaphases (Fig 3). In total, counts of germline-specific (LC positive) and somatically retained chromosome yield a haploid number of n=96 chromosomes in the lamprey germline, which is fewer than previously estimated n=99 [18, 34].

**Fig 3.**
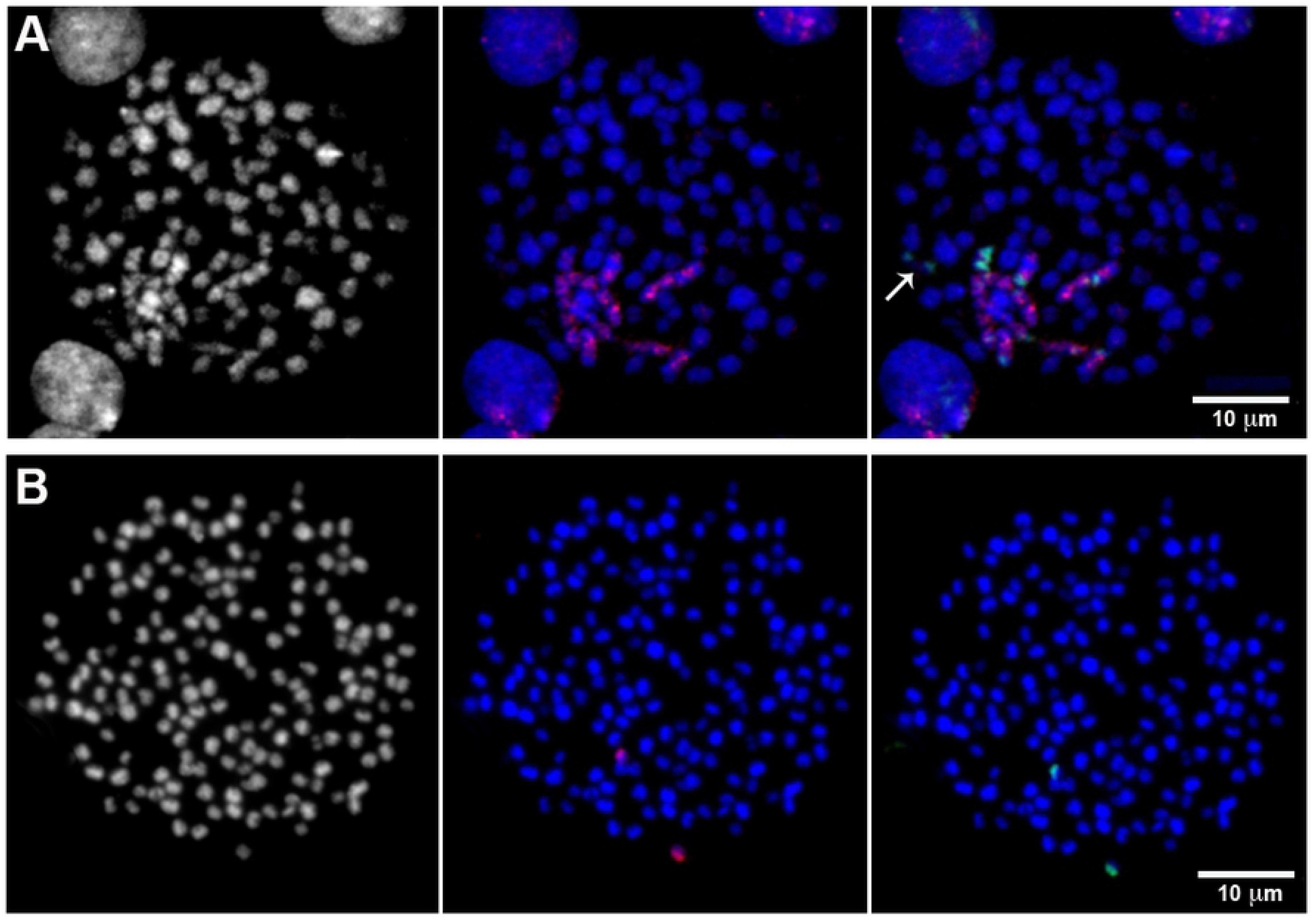
FISH of a probe generated from laser capture of lagging chromatin (red) and *Germ1*-repeat (green) on meiotic and somatic mitotic chromosomes. (A) hybridization to meiotic chromosomes from testes. The arrow on panel A indicates bivalent that corresponds somatically retained *Germ*1 enriched chromosome visible in somatic spreads. (B) A somatic mitotic metaphase.

In an attempt to uniquely identify individual germline-specific chromosomes, our six new computationally-predicted germline-specific repetitive elements (*Germ*2-7) and *Germ*1 were hybridized to metaphase chromosomes in three successive rounds of hybridization, which allowed us to localize all seven elements and the germline LC probe on the same set of meiotic metaphases. We analyzed at least 40 meiotic metaphase spreads and found that all 6 of the predicted germline-specific repeats hybridized exclusively to chromosomes that were marked by the germline LC probe (Fig 4). These hybridization patterns allowed us to verify that these elements are restricted to eliminated chromosomes and provided a means of individually identifying all 12 eliminated chromosomes.

**Fig 4.**
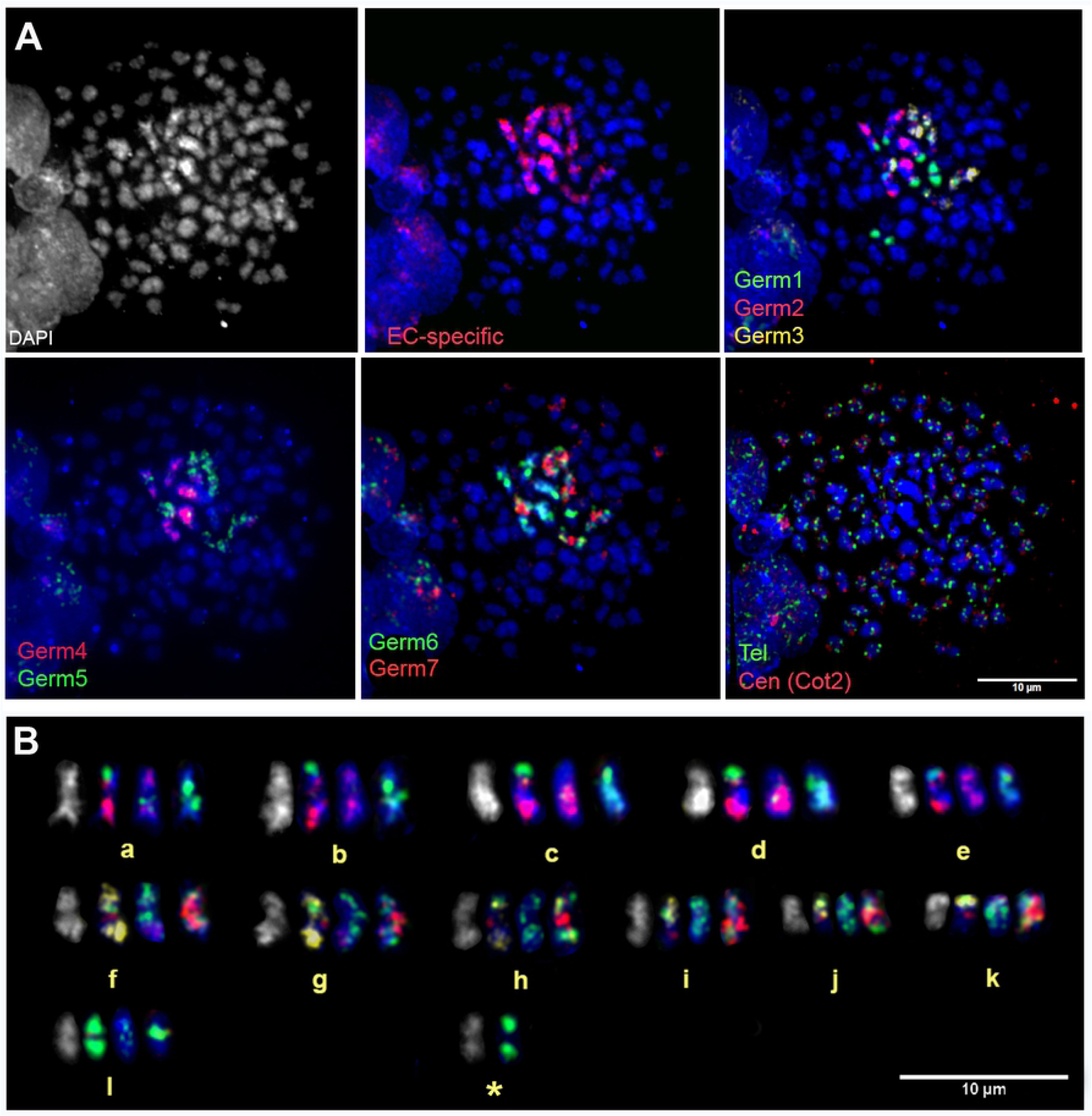
Chromosomal localization of the germline restricted repeats. (A) FISH of *Germ*1-7, repeats and telomeric/centromeric (liver C_0_t2) probes on spermatid spreads, with DAPI counterstaining (grey or blue). (B) A karyogram of germline restricted chromosomes (including a somatically-retained bivalent that cross-hybridized with *Germ1*: labeled with an asterisk). This bivalent presumably encodes somatic ribosomal RNAs, which cross-hybridize with *Germ*1 [18].

These hybridizations were used, in conjunction with reverse DAPI staining patterns, to develop an idiogram for all germline-restricted chromosomes and a map of the locations of *Germ*1-7 repeats on each chromosome (Fig 5). Remarkably, three repeats (*Germ* 1, 2, and 6) were found to be present on each of the 12 eliminated chromosomes, albeit with distinct distributions. *Germ*1 is present as dense signals that are located adjacent to the pericentromeric regions of all 12 eliminated chromosomes. This pattern contrasts with those of *Germ*2, which is typically located closer to the telomere, and *Germ*6 which generally shows a more diffuse patterns across the length of chromosomal arms. Unlike *Germ*1, the elements *Germ*2 and *Germ*6 do not cross hybridize with somatic chromosomes. The other four germline-specific repeats vary more broadly in their distributions across chromosomes and individual elements appear to be completely absent from one or more chromosome (Fig 4).

**Fig 5.**
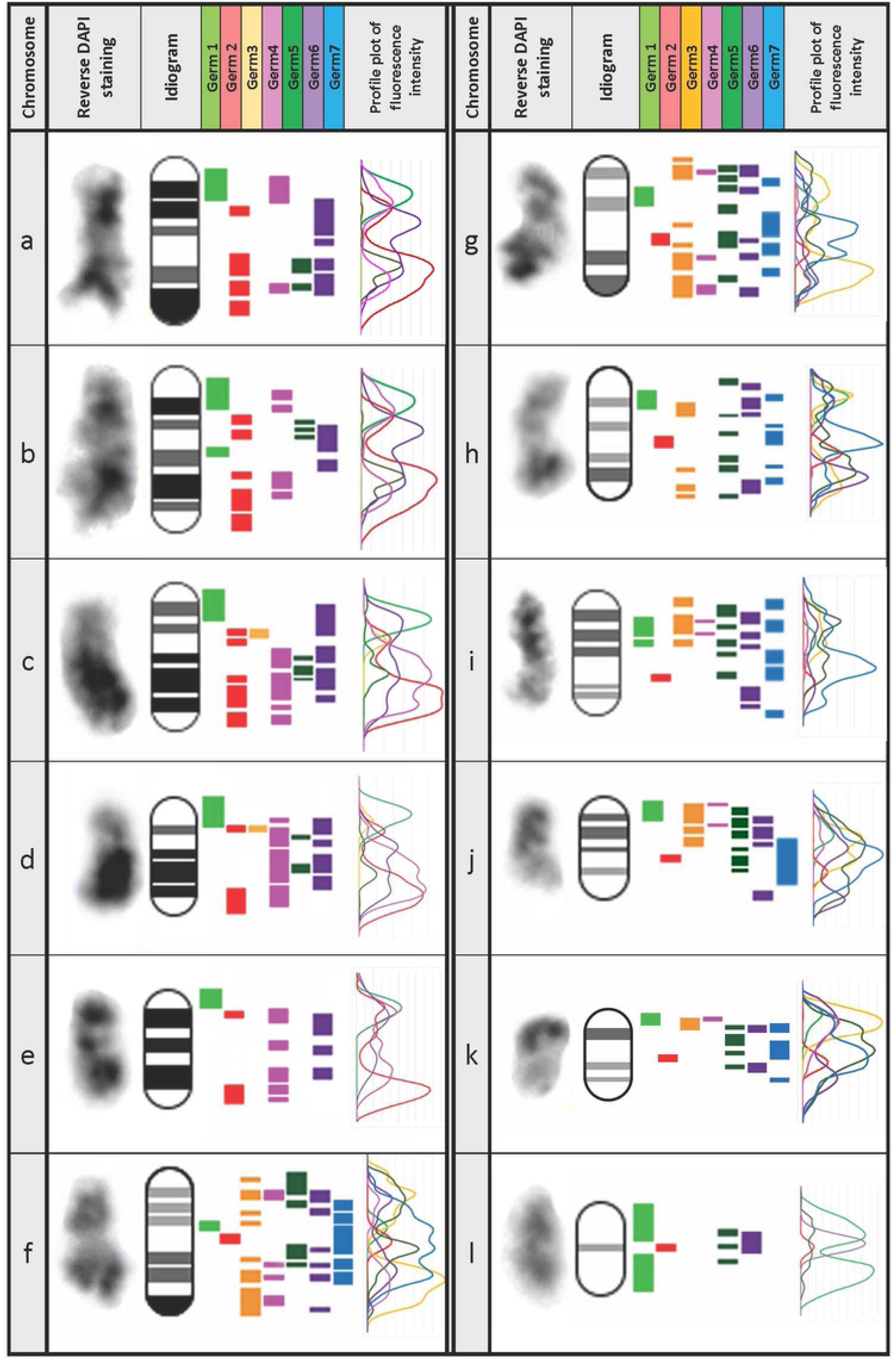
An idiogram of germline-specific chromosomes. Twelve germline-specific chromosomes (a-l) can be distinguished by their DAPI staining and hybridization patterns of *Germ*1-7 repetitive elements in meiotic metaphase-I spreads (Fig 4). Profile plots were generated based on the fluorescence intensity of hybridized DNA-probes corresponding to each repeat.

Over the course of examining metaphase spreads it was noted that germline-specific chromosomes were generally clustered within meiotic spreads. Closer examination of these clusters revealed that groups of germline-specific chromosomes frequently contacted one another near their telomeres, forming structures reminiscent of meiotic chains (Fig 6). Similar meiotic chains or ring formations have been described in evening primrose [37], *Incisitermes schwarzi* termites [38], *Leptodactylus* frogs [39] and platypus [40]. In these species, the formation of multivalent chains is thought to be associated with chromosomal translocations, as has been observed in mice with the Robertsonian exchange [41, 42]. However, in the sea lamprey meioses the number and content of chromosomes in each chain appears to vary from cell to cell (S4 File). We speculate that the formation of these variable chains during meiosis might reflect a general tendency for germline chromosomes to interact with one another at their distal ends, foreshadowing stronger interactions that occur during elimination (see below).

**Fig 6.**
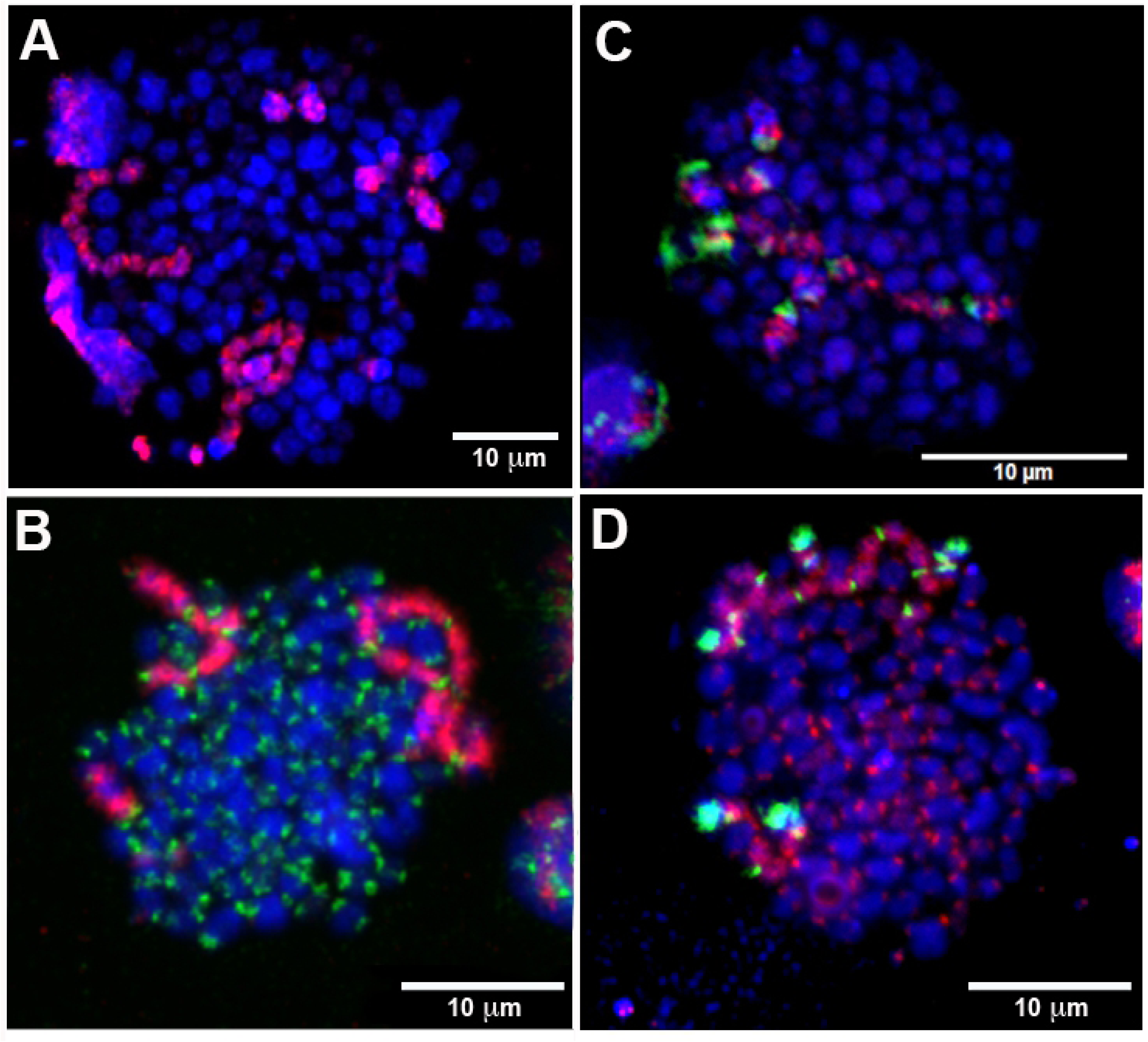
Meiotic chains in sea lamprey spermatid metaphase spreads. Four different meiotic metaphase-I spreads are shown. (A) A spread hybridized with the germline-specific LC probe (red) (B) A spread hybridized with the germline-specific LC probe (red) and liver C_0_t2 (green). Liver C_0_t2 DNA marks the location of centromeric repeats. (C) A spread hybridized with the germline-specific LC probe (red) and repeat *Germ*1 (green). (D) A spread hybridized with the germline-specific LC probe (red) and repeat *Germ*2 (green). All chromosomes are counterstained with DAPI (blue).

### Chromosomal Localization of Germline-Restricted Repeats in Elimination Anaphases

In order to directly resolve the spatial organization of germline-specific repeats during elimination, we performed 3D *in situ* hybridizations within fixed and PACT-cleared embryos (Fig 7). As expected from analyses of meiotic spreads, *Germ*2-7 signals were visible only in lagging (eliminated) chromatin and were absent from retained somatic chromosomes (Fig 4 and 7). Each of these repeats marks a distinct subregion of eliminated chromatin that corresponds to its relative position in meiotic metaphases, with centromeric ends (usually exhibiting *Germ*1 signals) being oriented toward the spindle poles. These signals are generally oriented in an antiparallel pattern along to the axis of anaphase elongation (suggesting telomeric/subtelomeric interactions between sister chromatids), with distal portions of the eliminated chromosomes remaining near the metaphase equatorial plane and proximal ends oriented toward the somatically retained chromosomes.

**Fig 7.**
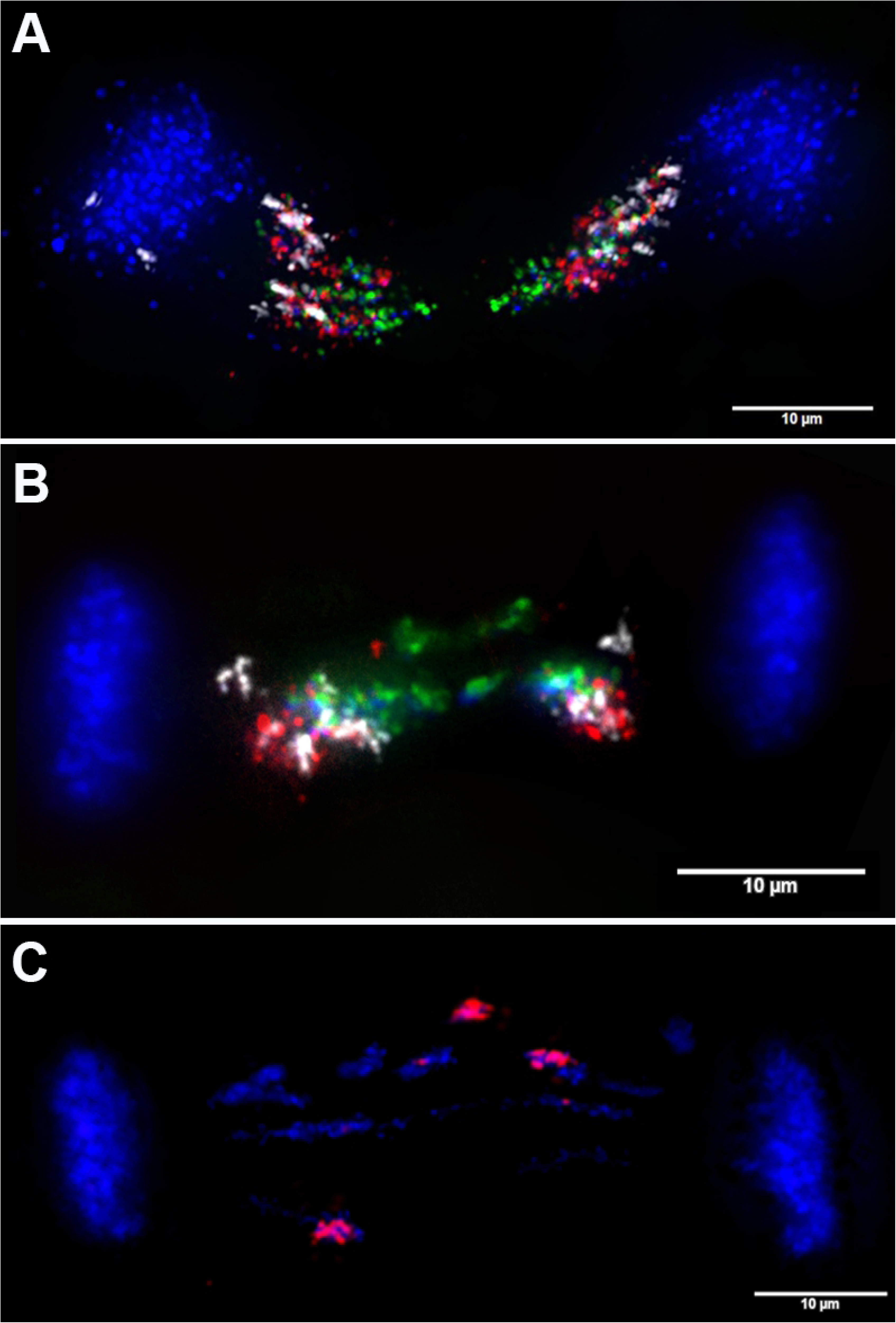
Lagging chromosomes and germline-restricted repeats. FISH of seven repetitive elements to lagging anaphases from 1.5 dpf sea lamprey embryos: (A) *Germ1* (white), *Germ2* (green), *Germ6* (red); (B) *Germ5* (red), *Germ4* (green), *Germ3* (white); (C) *Germ7* (red).

*Germ*2 and *Germ*4 elements were found frequently localized to the midline of lagging anaphases (Fig 7 A, B) suggesting the possibility that these repeats may mark a domain of interaction between the telomeres or subtelomeric regions of several chromosomes that possess dense *Germ*2 and *Germ*4 domains near their distal telomeres (Fig 5). To examine whether stretched lagging chromosomes retain intact telomeres on both arms, we carried out FISH using PNA (Peptide Nucleic Acid) probes to the vertebrate telomere repeat consensus sequence. These hybridizations reveal that stretched lagging chromatin possesses telomere signals on both ends, consistent with the idea eliminated DNA is composed largely of entire germline-specific chromosomes (Fig 8 and S5 File). Moreover, it appears that poleward oriented (centromeric) signals are significantly larger and brighter than their equatorially-oriented counterparts. It is tempting to speculate, that lagging chromosomes are establishing telomeric contacts due to telomere shortening and the formation of adhesive ends. However, two observations speak against this simplistic interpretation. First, similarly variable hybridization intensities also observed in meiotic and mitotic metaphases (*i.e.* non-eliminating divisions; S3 File, S4 File E-I). Second, medially oriented telomere signals often appear as distinct pairs of signals within regions that are overlain by broader hybridization signals from *Germ*2 (Fig 8 and S5 File F). These observations suggest that interactions between lagging sister chromosomes may involve repetitive (*e.g. Germ*2, 4) or other sequences near the telomere ends, and perhaps not telomere fusion *per se*.

**Fig 8.**
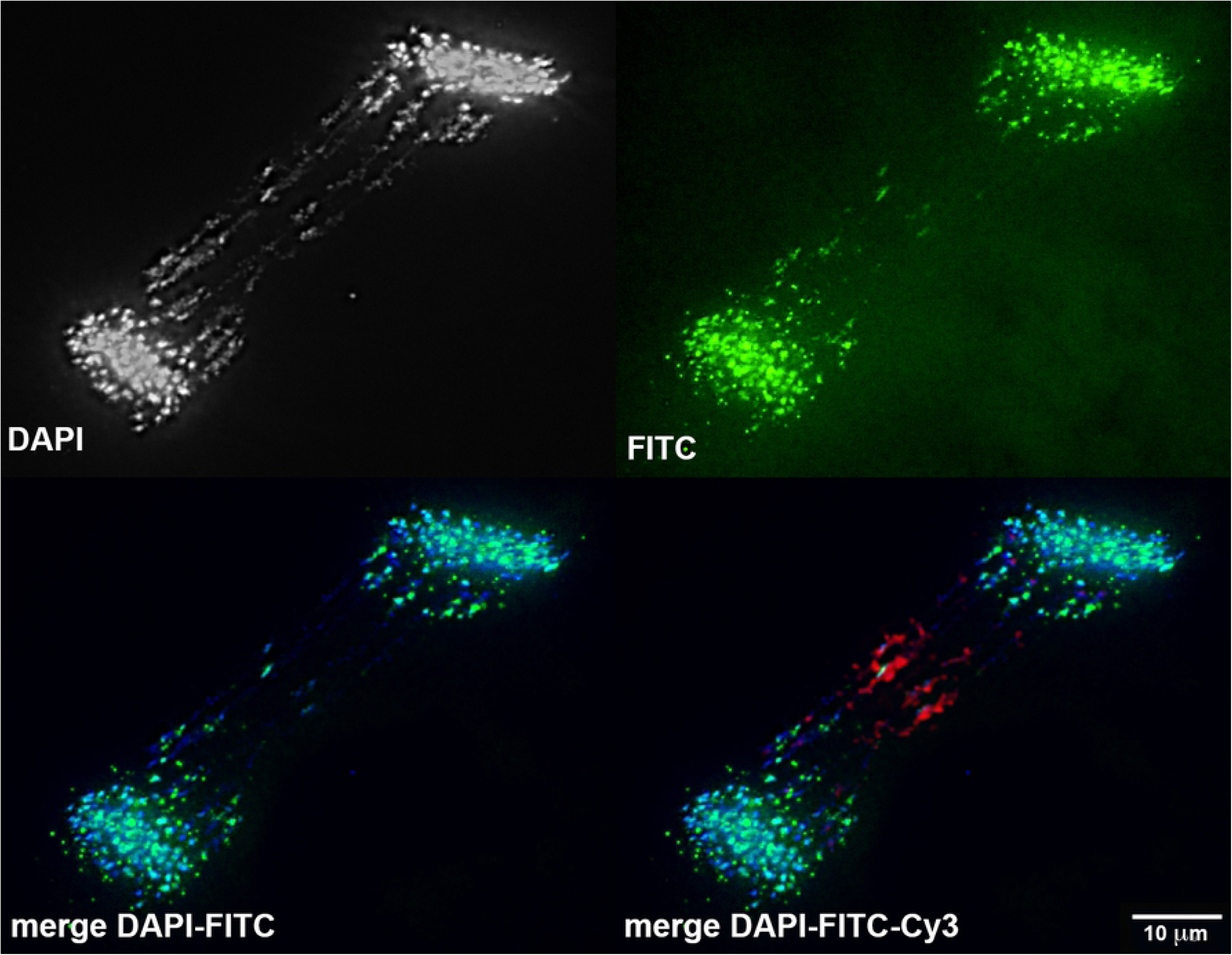
Telomeric contacts in chromosome elimination of the sea lamprey. Fluorescence *in situ* hybridization telomere specific (green) and Germ2 (red) probes on anaphase 1.5 dpf embryo (see also S5 File).

The identification germline-specific repetitive elements and localization of these elements to meiotic and eliminating cells sheds new light on the number and structure of eliminated chromosomes, as well as their behavior and sub-cellular orientation during elimination. Taken together, our observations suggest that interactions between lagging chromosomes result in the loss of large segments of the genome and are consistent with the loss of twelve entire germline-specific chromosomes. Analyses of the distribution of repetitive sequences across micronuclei (wherein lagging material is packaged prior to degradation) further indicate that some fragmentation likely takes place during the early stages of elimination, resulting in the formation of micronuclei that contain germline-specific DNA, but lack *Germ*1 (*Germ*1 cassettes are present on all eliminated chromosomes described here). This builds on earlier work demonstrating that early interactions and packaging events are followed by the accumulation of methylation marks and degradation of eliminated DNA within micronuclei [2] (Fig 9).

**Fig 9.**
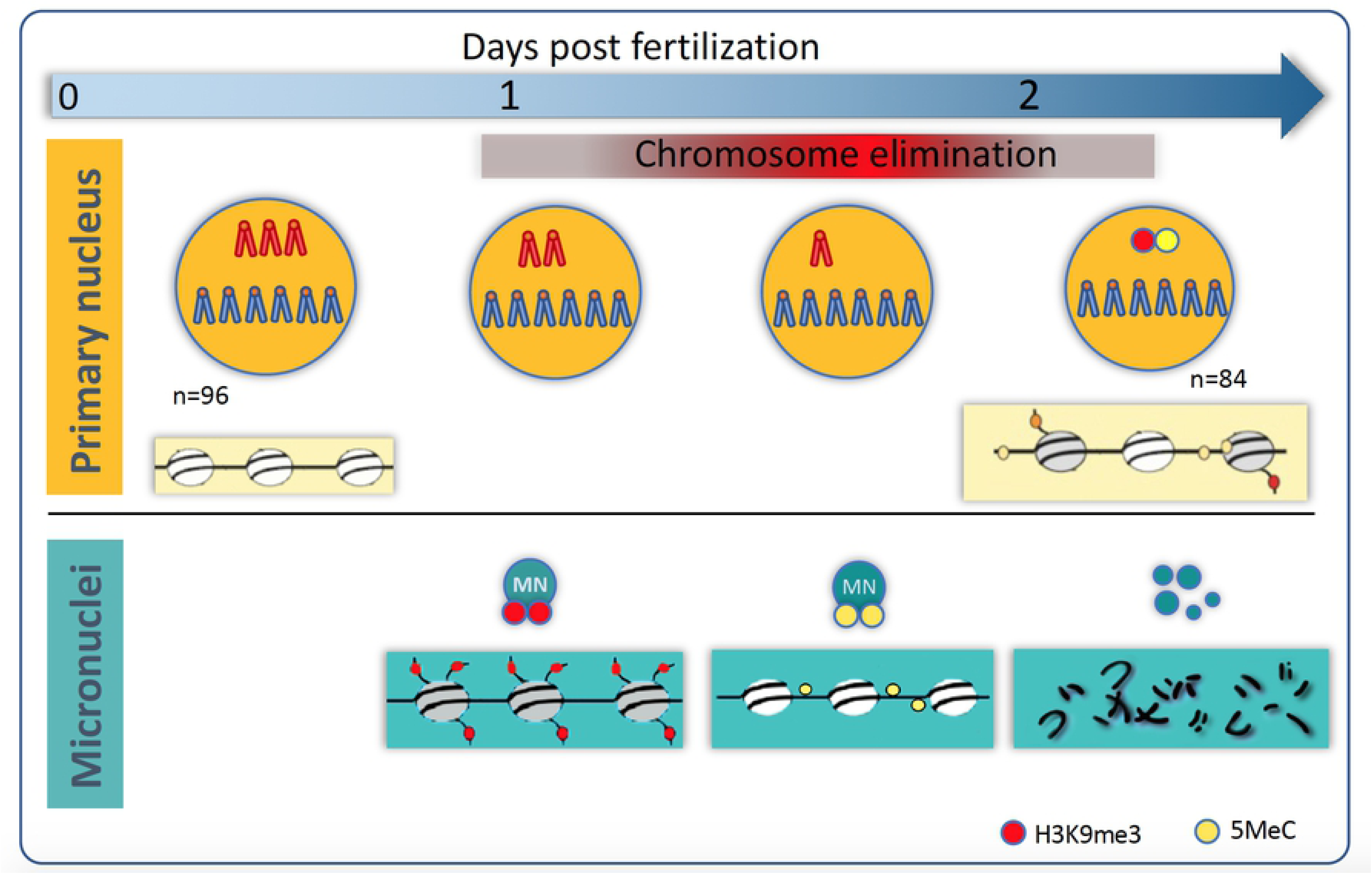
Chromosome elimination during the sea lamprey development. The initial haploid chromosome number in embryonic cells is n=96. Chromosome elimination is initiated after the 6^th^ cleavage at ∼1 dpf. Presomatic cells sequester 12 chromosome pairs into micronuclei that are initially enriched with silencing histone modifications. These histone modifications are subsequently replaced with 5-methylcytosine, which presumably reinforces transcriptional silencing. Primary nuclei show signs of epigenetic silencing only after the formation and remodeling eliminated chromosomes, at a time point that is roughly concurrent with the *in situ* degradation of germline-specific DNA.

These analyses highlight the highly regulated nature of PGR in lamprey and suggest an essential role of germline specific repetitive elements in targeting large DNA segments (chromosomes) for elimination. We anticipate that further analyses of these repeats and their associated binding partners in eliminating cells will provide new insights into mechanisms that mediate chromosome segregation and the maintenance of genetic integrity during mitosis, and might ultimately be coopted as a means of experimentally manipulating the behavior of specific intact chromosomes during anaphase. Our analyses demonstrate that lagging chromosomes likely have all fundamental morphological attributes necessary for normal chromosome segregation: their centromere ends orient toward the spindle poles, the chromatin itself situated along the spindle filaments implying interactions with spindle motor proteins [2], and lagging chromosomes retain intact telomeres. Moreover, they appear to segregate normally during the first several embryonic cell divisions and in the germline. We anticipate that further work aimed at understanding factors that interact with germline-specific sequences (including repetitive elements) will shed light on the cellular mechanisms that identify germline-specific sequences and mediate their differential motion during anaphase.

## MATERIALS AND METHODS

### Research animals

Animals were obtained from Lake Michigan via the Great Lakes Fisheries Commission and maintained under University of Kentucky IACUC protocol number 2011-0848 (University of Kentucky Institutional Animal Care and Use Committee). For tissue sampling, animals were euthanized by immersion in buffered tricaine solution (1.0 g/l), dissected, and tissues for DNA isolation were immediately frozen. For meiotic chromosome preparations, testes were extracted from non-spermating males, collected temporally in 1xPBS before processing.

### Production of lamprey embryos

*In vitro* fertilizations were performed with sexually mature adult animals. Eggs and sperm were collected in crystallization dishes and allowed to incubate in 10% Holtfreter’s solution for 10 min to permit fertilization [43]. After visually confirming activation, embryos were rinsed in distilled water to remove excess sperm and maintained in 10% Holtfreter’s solution at 18°C throughout development [44]. At days 1, 1.5, and 2 post-fertilization, live embryos were collected in 15 ml centrifuge tubes and fixed were fixed in MEMFA Fixative for 1 hour, rinsed in 1X PBS, dehydrated in increasing concentrations of methanol, and stored in methanol at −20°C as previously described [45].

### PACT clearing

MEMFA fixed embryos were embedded in hydrogel and cleared according to a PACT (passive clarity technique) protocol optimized for lamprey embryos [35, 46]. Prior to clearing, embryos were gradually rehydrated in 1X PBS then perfused with hydrogel monomer solution (5% acrylamide supplemented with 0.5% VA-044) by incubating overnight at 4°C. Hydrogel polymerization was performed at 37°C for 2.5 hours. After brief washes with 1X PBS, embryos were transferred to 50 ml screw-cup tube and incubated in stripping solution (8%SDS in 1X PBS) for 5 days at 37°C with gentle rotation. Upon reaching transparency samples were washed in 1X PBS with 5 buffer changes over the course of a day and transferred into staining solution (1X PBS, pH=7.5, 0.1 Triton X-100, 0.01% sodium azide). Cleared embryos were stored at room temperature prior to downstream processing.

### Preparation of metaphase spreads

Meiotic cell preparations were made from testes of non-spermating males. About 1 cm^3^ of tissue was homogenized in Dounce Grinder, the cells were treated with HEPES (0.01 M) buffered 0.075 M KCl hypotonic solution, pH=7.4 and fixed with methanol:glacial acetic acid (3:2) fixative solution. Mitotic spreads were obtained from 16 dpf embryos using overnight exposure to colchicine (0.04%), homogenization, and buffered hypotonic treatment. Cell suspensions were applied to a clean steamed glass slide and immediately placed in a humidity chamber at 55°C to facilitate proper chromosome spreading [47].

### C_0_t DNA isolation

For highly repetitive DNA (C_0_t) isolation, genomic lamprey DNA was extracted by phenol-chloroform method [48] and repetitive fractions were isolated using S1 nuclease to digest single stranded (low copy) DNA as described previously [49].

### Computational prediction of germline-specific repeats

Abundant k-mers (k = 31) were identified from lamprey sperm (SRR5535435) and blood (SRR5535434) DNAseq datasets using Jellyfish version 2.2.4 [50]. Minimal copy-number thresholds for defining abundant k-mers were set at 3X the modal copy number: 165 for sperm and 180 for blood. Abundant k-mers were extracted and assembled into a set of high-identity repetitive elements using Velvet version 1.2.10 [51] with a hash length of 29.

These *de novo* assembled repeats were aligned to repetitive sequences generated from the lamprey reference genome (PIZI00000000.1) by RepeatModeler [36]. Sequences aligning to RepeatModeler repeats with >90% identity over 80% of their length were replaced by the corresponding longer sequence. The previously-characterized *Germ*1 repeat was also added to the set leading to the exclusion of 13 repeats that matched it with 99% identity.

An enrichment analysis was performed by separately aligning paired-end reads from sperm (SRR5535435) and blood (SRR5535434) DNAseq datasets to the set of assembled reference repeats using BWA MEM. The DifCover pipeline [52] was used to calculate enrichment scores and stage 2 of the analysis pipeline was run with parameters that 1) prevent splitting sequences to intervals shorter than 5Kb (v=5000); 2) report intervals of any length (l=0); 3) assure reliable minimal coverage (a=b=10) by either sperm or blood reads; and 4) allow bases with depth of coverage as high as the observed maximum of 11.8M (A=B=12M). A set of 118,634 intervals generated by DifCover was filtered to identify 171 highly abundant and germline-specific sequences with enrichment scores of more than 5 and estimated span size of more than 40 Kb (S1 and S2 Tables). The estimated genomic span of these repeats was computed as [length of sequence*(sperm coverage/modal sperm coverage], where modal sperm coverage = 73.

Clustering of 171 highly abundant and germline-specific sequences was performed using CD-HIT-EST (v4.6, with paramaters: -c0.8, -G0, -aS 0.3, -aL 0.3, -sc 1, -g 1, -b 4) [53], resulting in the identification of 30 clusters. We then cross aligned (blastn with -word_size 11) [54] sequences from separate clusters and found that some clusters could be further merged (required to have at least 4 hits), resulting in 20 clusters, 8 of which contained multiple sequences and 12 of which were singletons. For characterization of larger-scale repetitive structures, genomic scaffolds with the largest number of hits to each germline-specific element were identified by BLAST alignment (blastn, -word_size 11, at least 80% of bases aligned) (S3 Table).

### Fluorescence in situ hybridization

#### Probes

To produce germline-specific probes, we used a DNA library that was generated from laser capture microdissected lagging anaphases as part of a previous study [19]. An aliquot of this library was amplified using the GenomePlex® WGA Reamplification Kit (Sigma) following the manufacturer instructions. Fluorescent LC probes were generated using a modified version of the manufacturer’s protocol: Cyanine 3-dUTP (Enzo), ChromaTide® Alexa Fluor® 594-5-dUTP (Thermo), or ChromaTide® Alexa Fluor® 488-5-dUTP (Thermo) were used along with 10 mM dATP, dCTP, dGTP, and 3 mM dTTP, replacing dNTP manufacturer kit mix. Probes for the *Germ*1 repeat and C_0_t1-2 fraction of genomic DNA were produced using nick-translation of either an isolated BAC clone (*Germ1*) [18] or C_0_t fractions according to previously published protocol [48, 55] using Cyanine 3-dUTP (Enzo), Cyanine 5-dUTP (Enzo), or Fluorescein-12-dUTP (Thermo) as labeled nucleotides. The TelG-FAM PNA-probe (TTAGGG repeats, PNA Bio) was used as telomere specific probe. Probes for repetitive sequences were labeled via conventional PCR using a dNTP mixture containing 1 mM dATP, dCTP, dGTP, and 0.3 mM dTTP, and one fluorophore: Cyanine 3-dUTP (Enzo), Cyanine 5-dUTP (Enzo), Fluorescein-12-dUTP (Thermo), or ChromaTide® Alexa Fluor® 488-5-dUTP (Thermo). Each PCR amplification was performed using 1 μg of genomic DNA template, 30-34 PCR cycles and a 30 second extension step in order to obtain appropriately sized probes for FISH.

#### DNA in situ hybridization on whole embryos and cytogenetic slides

3-D FISH on whole embryos was performed according to a previously described protocol [2, 19, 56]. For individual hybridization experiments 4-5 embryos were placed in 2 ml tube and incubated in 8% sodium thiocyanate solution, overnight at 37°C, washed in 1xPBS for 1 hour, and placed in 50% formamide in 2xSSC for 2-3 hours at 45°C. For hybridization, 50% formamide/2xSSC solution was replaced with 30 μl hybridization mix consisting of 50% formamide, 10% dextran sulfate, 0.01% sodium azide, and 250 ng of labeled DNA-probe. Embryos were pre-incubated overnight at 37°C to permit penetration of probes, after which probe and target DNA was denatured by heating samples to 75°C for 5 minutes, chilled on ice for 2 min, and then moved to 37°C for overnight hybridization. Three subsequent washes: 50% formamide in 2xSSC, 0.4x SSC, 0.3% Nonidet-P40 and 2xSSC, 0.1% Nonidet-P40 for 15 min each at 45°C, were performed the following day. Embryos were placed on a slide in a drop of ProLong™ Gold Antifade Mountant with DAPI (ThermoFisher) and enclosed under a coverslip with slight pressure.

FISH on chromosome preparations was carried out according to a standard protocol for chromosome spreads [57] with modifications [49]. For hybridizations involving the LC probe, a 20x excess of unlabeled C_0_t2 DNA used to suppress hybridization to somatic repeats. After denaturation, each hybridization mixture was incubated at 37°C for 30 min, then applied to chromosome spreads that had been already denatured and dehydrogenized [55]. Probes that were co-hybridized with LC, were denatured separately and pooled with the LC probe after suppressive incubation, but before applying to the slides.

#### C_0_t2-CGH

Repetitive fractions of genomic DNA were isolated from testes and liver DNA that were extracted from an adult male via phenol chloroform purification, as previously described [48]. A total of 1.5 μg testes C_0_t2 DNA was labeled with Cy3-dUTP (Enzo) by nick-translation in a final volume of 50 μl; the same amount of liver C_0_t2 was labeled with fluorescein-12-dUTP (ThermoFisher). After labeling, probes were precipitated by adjusting the solution to 70% ethanol in presence of 20 μg single stranded sheared salmon sperm DNA (Sigma-Aldrich), followed by centrifugation at 14,100 G. The resulting pellet was air dried and resuspended in 50 μl of hybridization buffer. 10 μl of each probe and 10 μl of hybridization buffer were mixed prior to hybridization to whole embryos. Conditions for hybridization were as described above.

#### Microscopy and image analysis

After FISH and immunostaining, slides were analyzed with an Olympus-BX53 microscope using filter sets for DAPI, FITC, TexasRed, and Cy5. Images were captured using CellSence software (Olympus). For thicker samples, such as intact cells from PACT cleared embryos, the extended focal imaging function was used in order to generate a single deep-focus image. Images from each filter set were captured separately, and merging of channels was performed using Adobe Photoshop CC 2017. Measurements of fluorescence intensity were carried out in ImageJ 1.48v (NIH) using raw images and options “integrated density” and “mean gray value” for selected area on captured images.

## SUPPORTING INFORMATION

**S1 File. C_0_t CGH on anaphase chromosomes of the sea lamprey embryos undergoing germline chromosome elimination**. Germline repetitive DNA (C_0_t2 isolated from testes) labels lagging chromosomes with equal or greater intensity in comparison to somatically retained chromosomes, whereas the fluorescence produced by a liver C_0_t2 DNA probe is almost completely absent from lagging chromosomes but well represented in somatically retained chromosomes.

**S2 File. Validation of germline enriched repeats via PCR.**

DNA extracted from testes ‘T’ was used as a germline sample, and DNA extracted from female muscle tissue as a somatic sample with reduced potential for germline contamination ‘fM’ [58]. Panels A-D show different levels of specificity of PCR reactions depending on the primer annealing temperature: (A) 65°C, (B) 60 °C, (C) 55 °C, (D) An additional 50°C PCR gel was run for repeat13 (not used in further analysis), Germ6, and Germ4.

**S3 File. FISH of centromeric (C_0_t1 DNA, *red*) and telomeric (*green*) probes on a somatic spread.**

Some chromosomes have clear difference in the strength of the telomeric signal at their ends.

**S4 File. Meiotic multivalents in the sea lamprey.**

Meiotic chains in sea lamprey metaphase I spreads (MI) hybridized with the LC probe in combination with various other probes. (A) LC probe only (red); (B – D) LC (red) and *Germ*1 (green); (E, F) LC (red), telomere PNA probe (green); (G) LC (red), *Germ*1 (green), telomere PNA probe (yellow); (H, I) left: LC (red), testes C_0_t2 (red), *Germ*1 (green); right: testes C_0_t2 (red), telomere PNA probe (green); (J-L) LC (red), liver C_0_t2 (green); (M, N) LC (red), genomic liver (green); Spreads A, B, I and L are also presented in Fig 6.

**S5 File. Telomere dynamics in lagging anaphases.**

Fluorescence *in situ* hybridization of the *Germ*2 probe (red), a PNA-probe for telomeric repeats (green, shown also in an additional panel), and testes genomic DNA (Cy5 labeled, pseudocolored in blue and gray-scaled in an additional panel) to eliminating anaphases of 1.5 dpf sea lamprey embryos. Stretched chromosomes typically have telomeric signals visible on both ends, equatorial telomeric signals appear as both merged (A, B, C, D, E) and split (C, F) signals.

**S1 Table. Enrichment scores (shown in descending order), estimates of the total genomic span and germline enrichment of repetitive sequences in the lamprey genome for intervals computed by DifCover within all modal repetitive elements.**

**S2 Table. Clustering of high-copy elements with enrichment scores exceeding 5 and estimated span exceeding 40 kb.**

**S3 Table. Repeats GERM2-7: enrichment scores, estimates of the total genomic span, sequence structure and PCR primers used for validation and generation of FISH probes.**

